# Subcortical rather than cortical sources of the frequency-following response (FFR) relate to speech-in-noise perception

**DOI:** 10.1101/2020.03.29.014233

**Authors:** Gavin M. Bidelman, Sara Momtaz

## Abstract

Scalp-recorded frequency-following responses (FFRs) reflect a mixture of phase-locked activity across the auditory pathway. FFRs have been widely used as a neural barometer of complex listening skills, especially speech-in noise (SIN) perception. Applying individually optimized source reconstruction to speech-FFRs recorded via EEG (FFR_EEG_), we assessed the relative contributions of subcortical [auditory nerve (AN), brainstem/midbrain (BS)] and cortical [bilateral primary auditory cortex, PAC] source generators with the aim of identifying which source(s) drive the brain-behavior relation between FFRs and SIN listening skills. We found FFR strength declined precipitously from AN to PAC, consistent with diminishing phase-locking along the ascending auditory neuroaxis. FFRs to the speech fundamental (F0) were robust to noise across sources, but were largest in subcortical sources (BS > AN > PAC). PAC FFRs were only weakly observed above the noise floor and only at the low pitch of speech (F0≈100 Hz). Brain-behavior regressions revealed (i) AN and BS FFRs were sufficient to describe listeners’ QuickSIN scores and (ii) contrary to neuromagnetic (MEG) FFRs, neither left nor right PAC FFR_EEG_ predicted SIN performance. Our preliminary findings suggest subcortical sources not only dominate the electrical FFR but also the link between speech-FFRs and SIN processing as observed in previous EEG studies.

## 1. INTRODUCTION

The frequency-following response (FFR) has provided considerable insight into how well auditory neural coding relates to perception, particularly speech-in-noise (SIN) listening skills. FFRs are neurophonic potentials generated by a mixture of subcortical and cortical structures along the auditory system that phase-lock to spectrotemporal features of periodic sounds including speech. Links between speech-FFRs (recorded with EEG) and SIN processing have been widely reported over the past decade [e.g., 9, 22, 23, 27, 33, 35]. For instance, Song, et al. [33] showed the magnitude of the speech fundamental frequency (F0) encoded via FFRs positively predicted SIN performance on the QuickSIN [15]: “Top SIN” performers on the task had more robust FFRs than “Bottom SIN” performers, who had both weaker neural representation of the speech F0 and poorer perceptual scores. Complementary findings were reported by Anderson, et al. [1] who showed poorer (i.e., lower median) SIN listeners experienced greater (∼0.5-1 ms) noise-related shifts in the timing of their speech FFR from quiet to noise than top performing listeners. These findings suggest a relationship between FFRs to complex sounds and SIN perception, whereby faster and more robust speech encoding is associated with better behavioral outcomes.

Though traditionally viewed as a brainstem potential [32], it has long been recognized there are multiple sources of FFRs stemming from throughout the hearing pathway. These include the cochlea [31], auditory nerve [5, 6], upper brainstem (midbrain inferior colliculus) [5, 6, 32], and under some circumstances, primary auditory cortex [6, 10]. EEG studies have interpreted FFR correlates of SIN perception based on how well *(brainstem)* FFRs encode important acoustic properties of speech (e.g., voice pitch and timbre cues) [9, 22, 23, 33]. Still, acknowledging FFRs likely contain contributions from cortex for low-frequency (∼100 Hz) stimuli [6, 10], it is possible the link between this neurophonic and SIN behaviors is at least partially driven by higher-level “cortical FFRs” [cf. 10, 11]. Based on MEG, it has been suggested higher-level cognitive tasks such as SIN perception were dominated by FFRs generated in right auditory cortex [11, see also 13]. This finding is in stark contrast to the brainstem-centric view of the FFR and its relation to auditory perception [17, 23, 27, 33, 36, 37].

We suspect ambiguity of cortical vs. subcortical structures in accounting for perceptual correlates via FFR might be driven by limitations of different neuroimaging modalities. Whereas subcortical structures are among the most significant sources of the scalp FFR_EEG_ [5, 6, 25], MEG is more sensitive to superficial neuronal activity [deep brainstem sources become invisible; 12, 17]. Given the differential pickup of deeper brainstem (EEG) vs. shallower cortical (MEG) contributions to the FFR, it is reasonable to assume that imaging modality could bias associations between FFR and behavior. Moreover, phantom simulation studies reveal MEG localization accuracy of deep sources is greater for gradiometer vs. magnetometer recordings [most MEG-FFR studies use the former; 10, 11, 25], suggesting the detection of neuromagnetic brainstem activity depends even on the choice of sensor itself [2]. Because FFRs are recorded more commonly with EEG, we were interested to reevaluate the neural origin(s) of the FFR_EEG_-SIN association to determine *which* subcortical and/or cortical structure(s) drives this relation.

In light of emerging controversies on sources of the FFR and their relation to complex auditory behaviors [6, 10], we aimed to unravel the link between speech-FFRs and SIN processing using a more comprehensive, systems-level neuroimaging approach. Using high-density EEG, we recorded multichannel FFRs to noise-degraded speech. Source analysis allowed us to parse region-specific activity underlying the electrical FFR and evaluate the relative contributions of each nuclei to SIN perception. Our findings show phase-locked activity peripheral to cortex dominates the EEG-based FFR as well as its link to perceptual speech-in-noise listening abilities.

## 2. METHODS

### 2.1 Participants

Data herein represent FFRs originally recorded in *n*=12 normal-hearing adults (age: 24.7±2.7 years) [7]. All participants were native English speakers with a similar level of education (undergraduate degree or higher) and <3 years formal music training (1.3±1.8 years) that occurred at least five years before the study. Hearing thresholds were normal (≤ 25 dBHL) bilaterally at octave frequencies between 250-8000 Hz in all participants. All gave written informed consent for the study protocol approved by the University of Memphis Institutional Review Board.

### 2.2 Stimuli

FFRs were elicited by a 300 ms male speech token /ama/ [for details, see 7]. The F0 pitch fell gradually over its duration (F0= 120-88 Hz). The low F0 of the stimulus (∼100 Hz) was expected to elicit phase-locked FFRs of both subcortical and cortical origin [6, 10]. In addition to this “clean” stimulus, speech was presented in continuous four-talker babble noise [15] at signal-to-noise ratios (SNRs) of +10 and +5 dB. Listeners heard 2000 trials of each speech token (passive listening) per noise block presented at 81 dB SPL through ER-30 insert earphones (Etymotic Research). Extended acoustic tubing (20 ft) with the headphone transducers placed outside the booth avoided electromagnetic stimulus artifact from contaminating FFRs.

### 2.3 EEG

EEG recording and preprocessing followed previous reports [7]. EEGs were digitized (5000 Hz sampling rate; online filters = DC-2500 Hz) from 64 electrodes at 10-10 scalp locations. EEGs were then ocular artifact corrected, epoched (−200-550 ms), baseline corrected, common average referenced, and ensemble averaged to obtain FFRs for each noise condition. Responses were bandpass filtered (80-1500 Hz) for subsequent source analysis.

#### 2.3.1 Source waveform derivations

Scalp FFRs (sensor-level recordings) were transformed to source space using a virtual source montage implemented in BESA® Research v7 (BESA, GmbH) [7]. This digital re-montaging applies a spatial filter to all electrodes (defined by the foci of our dipole configuration) to transform electrode recordings to a reduced set of source signals reflecting the neuronal current (in units nAm) as seen *within* each anatomical region of interest. We adopted the dipole configuration described in Bidelman [6] to assess the relative contribution of subcortical and cortical FFR sources to SIN processing. Full details of this model, its derivation, and fit accuracy are reported elsewhere [6]. The model consisted of 5 dipoles seeded in bilateral auditory cortex (PAC; source #1−2), the upper brainstem (midbrain inferior colliculus) (BS; source #3), and bilateral auditory nerve (AN; sources #4−5) (Fig. 2, inset). This allowed us to reduce listeners’ electrode recordings (64-channels) to 5 unmixed source waveforms describing their scalp FFR data. For each participant, dipole locations were held fixed and were used as a spatial filter to derive FFR source waveforms [6]. Critically, we fit individual dipole orientations to each participant’s data (anatomical locations remained fixed) to maximize the explained variance of the model at the individual subject level. The model provided a robust fit to the grand averaged (clean) scalp data across subjects in post-stimulus response interval (0-350 ms; goodness of fit: 77%^1^). Additional latency and spatial sensitivity analysis verified anatomical plausibility and good spatial separability of this dipole model (Fig. S1 and S2).

#### 2.3.2 FFR source waveform analysis

FFR source waveforms were analyzed in the spectral domain via BESA. From each waveform (per source, SNR, and listener), we first computed Fast Fourier Transforms (FFTs) in the post-stimulus response interval (0-350 ms, cos^2^ windowed; 2^11^ point FFT= 2.4 Hz resolution; see Fig. 1, shading). We then measured the peak amplitude in the F0 frequency bin. We focused on F0 because (i) this component was present across all sources of the FFR (see Fig. 2) and thus could be measured at each anatomical level and (ii) FFR-F0 has been explicitly related to SIN processing in previous EEG studies [7, 9, 24, 26, 29]. F0 amplitude was measured as the maximum spectral peak within the frequency range of 88­120 Hz [i.e., the F0 range of the stimulus pitch prosody; 7], relative to the noise floor [computed via the FFT of the pre-stimulus interval (−200-0ms)—interpolated to match the frequency resolution of the post-stimulus FFT]. Subtracting pre-stimulus from post-stimulus interval amplitudes thus retained only evoked neural responses (i.e., FFRs) above the EEG noise floor. Values below the noise floor where recorded as missing values (only 3 measurements, 2 of which were from PAC). This allowed us to assess the degree to which subcortical and cortical FFR sources showed veridical phase-locked activity (i.e., above noise-floor). For each source/SNR/participant, F0 measurements were scored twice and subsequently averaged. Test-retest reliability was excellent (*r* = 0.99).

**Figure 1:**
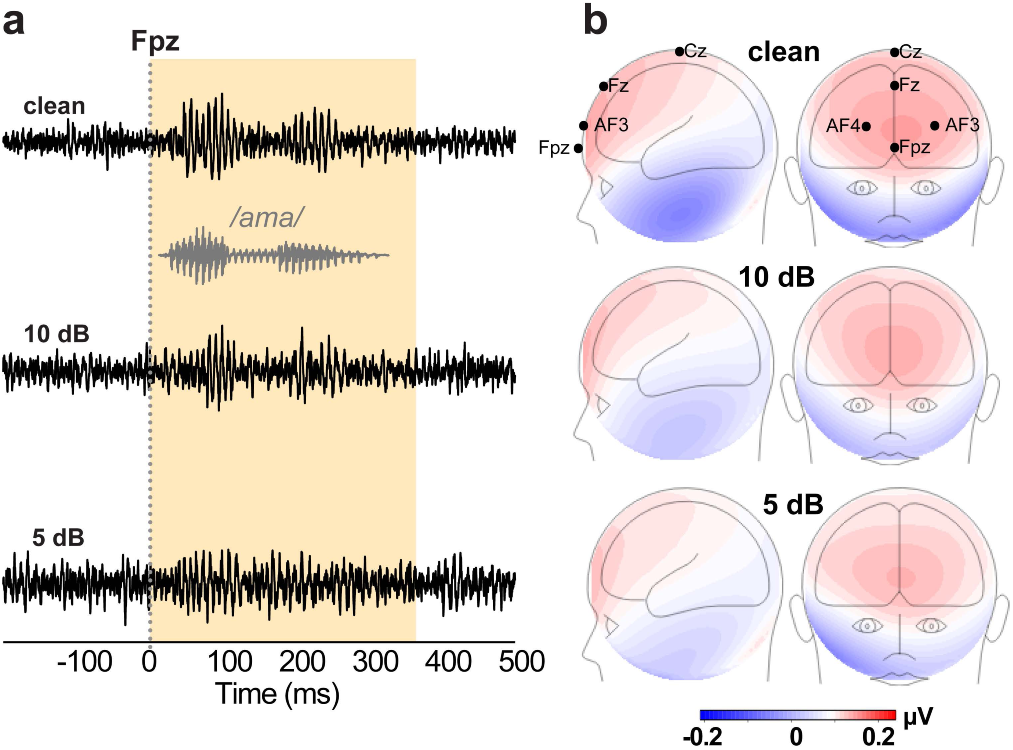
FFR (sensor-level) waveforms and scalp topographies as a function of noise. (**a**) Electrode recordings at channel Fpz across noise levels. Gray trace= stimulus waveform. FFRs appear as phase-locked potentials that mirror the acoustic periodicity of speech. (**b**) FFR topographies. Maps reflect the voltage distribution on the scalp, averaged across the periodic “wavelets” of the FFR [i.e., 25 most prominent positive peaks; see 5] in the response time window (0-350 ms; yellow shading). Maximal FFR amplitude near frontocentral sites (e.g., Fpz) and polarity inversion at the mastoids are consistent with deep midbrain sources that point obliquely in an anterior orientation to the vertex (parallel to the brainstem) [5, 6]. Red/blue shading = +/− voltage.

**Figure 2:**
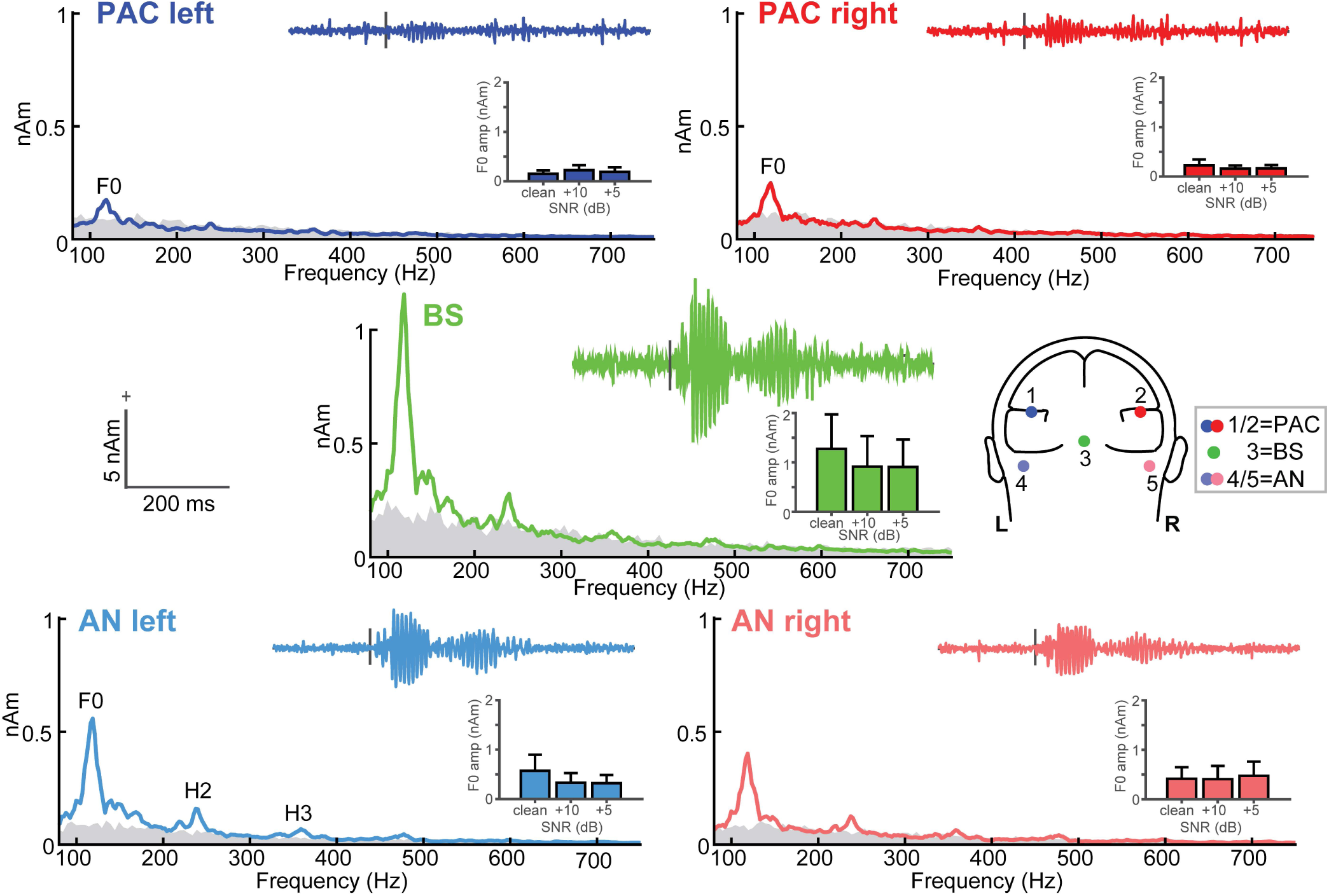
Source-level FFR waveforms and spectra along the ascending auditory neuroaxis. Grand average response spectra and waveforms *(insets)* at each dipole source of the FFR [head model; 6]. Waveforms and FFRs reflect responses to clean speech. Inset bar charts show noise-related changes in FFR F0. Gray shading = spectral noise floor measured in the pre-stimulus interval. FFRs show strong phase-locking at the speech F0 frequency (∼100 Hz) in both subcortical and cortical sources. Subcortical sources (AN, BS) show additional response energy at higher harmonics of speech (H2 and H3). errorbars = ±1 s.e.m.

### 2.4 QuickSIN task

Behavioral SIN reception thresholds we measured by QuickSIN [15]. Participants heard two lists of six sentences embedded in four-talker babble noise. Each contained five key words. Sentence presentation was 70 dB SPL amid noise that decreased in 5 dB steps from 25 dB (very easy) to 0 dB SNR (very difficult). “SNR loss” (in dB) was determined as the SNR required to correctly identify 50% key words [15]. Two lists were averaged to obtain a SIN score per listener^2^.

### 2.5 Statistical analysis

We used two-way, mixed-model ANOVAs (source x SNR; subjects=random factor) to assess changes in FFR F0 amplitude across anatomical levels and noise conditions. Amplitudes were SQRT-transformed to satisfy normality and homogeneity of variance. Generalized linear mixed effects (GLME) regression evaluated links between behavioral QuickSIN scores and the F0 component of source-localized FFRs [pooled across the noise (+10, +5 dB SNR) conditions]. Subjects were included as a random factor in the regression to model random intercepts per listener [e.g., QuickSIN ∼ PAC_left_ + PAC_right_ + BS + AN_left_ + AN_right_ + (1| subject)]. This multivariate model allowed us to assess the relative contribution of subcortical and cortical FFR sources to SIN perception.

We assessed reliability of the results using bootstrapping. Using the original dataset, participants were randomly resampled N=250 times (with replacement) in a leave one out scenario. From each resample, the GLME between QuickSIN scores and neural measures was recomputed and p-values for each source regressor were logged. Similarly, we used Bayes Factor analysis [19] to assess evidence in favor or against the null hypothesis. Bayes methods do not rely on large-sample theory and thus, are more appropriate for drawing inferences on smaller samples as used here [e.g., 21].

## 3. RESULTS

FFRs mirrored the periodicity of speech, reflecting robust phase-locking to the auditory input (Fig. 1). Their voltage distribution on the scalp showed maximal amplitude at frontocentral electrode sites (e.g., Fpz) and polarity inversion at the mastoids, consistent with sources in the deep midbrain that point obliquely in an anterior orientation to the vertex (parallel to the brainstem) [5, 6].

### 3.1 Source FFR waveforms and spectra

Source FFRs showed strong energy at the speech F0 frequency (∼100 Hz) in all source foci (Figure 2). However, only subcortical sources (AN, BS) showed response energy above the noise floor at higher harmonics of the speech signal (H2 and H3), consistent with the higher phase-locking limits of more peripheral auditory nuclei [14], including those generating the FFR [6]. In contrast, cortical PAC sources showed only weak energy at the F0 and failed to show reliable FFRs above the noise floor at higher frequencies^3^. Raw waveforms similarly show variation in the strength of the response across the neuroaxis. FFRs were maximal at the level of BS, were robust but to a lesser extent in AN, and reduced dramatically at the level of PAC.

### 3.2 FFR F0 amplitudes as a function of source level and noise

FFR F0 strength varied across source levels *(F*_4,126_*=* 10.42, *p* <0.0001) but not with SNR *(F*_2,126_*=* 0.47, *p* = 0.6286) (SNR × level: *F*_8,126_*=* 0.38, *p*=0.9313) (Fig. 2, *bar insets).* Tukey-Kramer comparisons revealed stronger F0 amplitudes in BS vs. PAC (*p*<0.0001), BS vs. AN (p=0.0002), and AN vs. PAC (p=0.0326). There was no difference between PAC_right_ vs. PAC_left_ (*p*=1.00). Subcortical sources were also collectively stronger than cortical sources (BS/AN > PAC; *p*<0.0001). Thus, our data did not reveal evidence of a functional asymmetry (i.e., right hemisphere bias) for cortical FFRs as observed in MEG studies [10]. Moreover, we found a midbrain-dominant gradient in F0 amplitudes across levels (BS > AN > PAC), further suggesting BS is the largest contributor to the FFR_EEG_ response [6].

### 3.3 Brain-behavior relations

Following previous channel-based FFR studies, we first evaluated associations between scalp-level recordings (Fpz electrode) and QuickSIN scores. We selected this channel as FFRs are strongest at frontocentral scalp locations (e.g., Fig. 1) [5, 6]. GLME regression revealed FFRs at Fpz were marginally predictive of better SIN perception (i.e., lower QuickSIN scores) (*F*_1,10_=4.75, *p*=0.054, adj-*R*^2^=0.69). Though non-significant here, these channel-level results parallel previous studies reporting links between FFRs and SIN processing observed in scalp-recorded data [e.g., 9, 22, 23, 33, 35].

We next adjudicated which FFR sources drive this brain-behavior relation and predict SIN performance. QuickSIN scores were uniformly good (0.38 ± 0.85 dB SNR loss), as expected for normal-hearing, but did span ±2 dB across listeners (Fig. 3a). GLME regression revealed the aggregate of FFR sources predicted QuickSIN performance [*F*_5,4_=7.95, *p*=0.0333; null hypothesis coefficients=0; adj-*R*^2^=0.87) (Fig. 3b). Bayes Factor analysis^4^ [19] indicated the alternative hypothesis was 3.72x more likely than the null hypothesis of no brain-behaver relation, suggesting moderate evidence for an FFR-QuickSIN relation [18]. Scrutiny of individual model terms revealed significant neural predictors in BS (β= −3.53; *p*=0.0071) and bilateral AN (AN_left_: β= −4.7, *p*=0.0062; AN_right_: β= −4.2, *p*=0.011). Contrary to MEG FFRs [11], neither left nor right PAC FFR_EEG_ predicted QuickSIN performance (PAC_left_: β= 0.67, *p*=0.70; PAC_right_: β= −2.18, *p*=0.13). Bootstrapping confirmed that among the five sources, only BS and AN were reliable predictors (*p*<0.05) among the surrogate datasets (Fig. 3c)^5^. Collectively, these findings suggest subcortical sources not only dominate the FFR_EEG_ but also the link between speech-FFRs and typical SIN performance [e.g., 9, 22, 23, 27, 33, 35].

**Figure 3:**
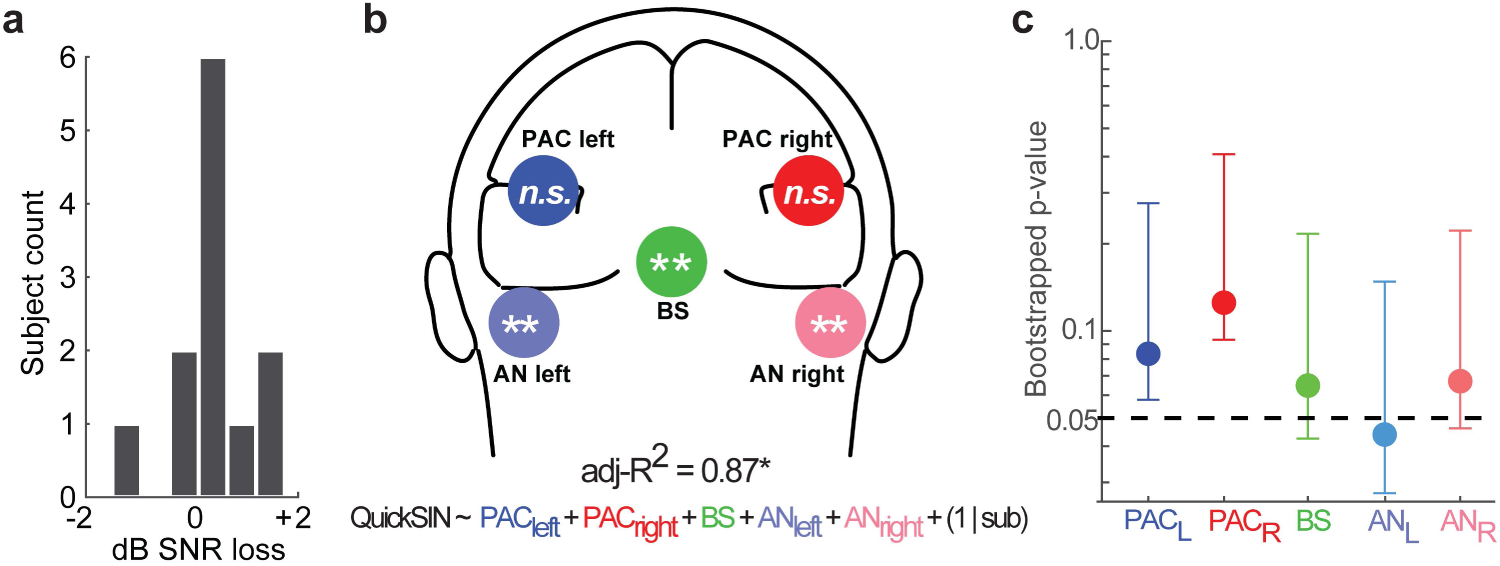
Subcortical speech coding is sufficient to explain SIN listening skills. (**a**) QuickSIN scores. **(b)** Subcortical and cortical sources of the FFR collectively account for 87% of the variance (adj-*R*^2^) in behavioral QuickSIN scores. Statistical flags mark significant predictors in the GLME model. Subcortical sources (left/right AN, midbrain BS) are associated with SIN performance, whereas cortical FFRs (left/right PAC) fail to predict behavior. (**c**) Bootstrapped p-values for each source regressor. Only BS and AN reach significance among N=250 resamples. Error bars = 95% CI. \**p* < 0.05; \*\**p* < 0.01

## 4. DISCUSSION

Our results confirm that while a mixture of auditory neural generators contribute to the aggregate FFR [6, 10], midbrain and lower structures dominate the FFR_EEG_ [e.g., 5, 6, 16, 30, 32, 34, 36]. Whereas subcortical (AN, BS) FFRs were robust, PAC showed comparably weak responses. In contrast, we find subcortical FFRs are not only larger at F0 than their cortical counterparts but persist at higher spectral frequencies to maintain coding of the harmonics and timbral elements of speech [6]. Our findings imply that previously observed relations between speech-FFRs and SIN perception [e.g., 7, 9, 22, 23, 27, 33] are mediated by subcortical sources.

### 4.1 Neural phase locking diminishes along the ascending auditory system

Our findings converge with human and animal studies suggesting subcortex (e.g., inferior colliculus and lower structures) as the primary source(s) of the FFR within the 20-200 Hz frequency bandwidth [6, 16, 30–32]. Diminishing FFRs from sub-to neo-cortex is consistent with the decreasing phase-locking tolerance of nuclei along the ascending auditory pathway [14] (**see** *SI Discussion*).

While our data show clear anatomically-dependent changes in FFR coding, we found noise had little impact on F0 responses. The lack of strong noise effects on pitch (F0) coding is consistent with prior FFR studies using relatively favorable SNRs (i.e., > 0 dB) [7, 9, 24, 26, 29]. Such resilience (and even increment) of F0 responses in the presence of mild-to-moderate noise is hypothesized to be due to stochastic resonance, the recruitment of low-frequency “tails” of high-frequency cochlear neurons due to high-intensity stimulus levels, and/or across channel integration of F0-related harmonics or intermodulation distortion products [for discussion, see 4]. All these putative mechanisms could lead to more robust neural synchronization and therefore redundancy of pitch-relevant cues carried in FFRs even with additive noise. Whether or not these factors affect each putative FFR_EEG_ generator remains to be tested.

### 4.2 Subcortical sources drive the link between speech-FFRs and SIN perception

Our data reveal SIN processing is governed by a coordinated orchestration of phase-locked activity to the spectrotemporal details of speech generated from different levels of the auditory system. However, our multivariate regression showed that subcortical sources (AN, BS) better predict listeners’ QuickSIN scores. In stark contrast, “cortical FFRs” failed to predict perception. Moreover, we find PAC responses are eradicated for the most meaningful frequencies in speech (i.e., energy above F0) which carry timbral information on talker identity, including important formant cues [9, 32]. This is not to say cortex is not involved in SIN perception. On the contrary, cortical ERPs suggest auditory and non-auditory brain regions in both hemispheres enable SIN perception [8]. Rather, our data reveal little-to-no involvement of cortex when it comes to the relation between sustained, stimulus phase-locked FFRs and SIN processing.

MEG studies suggest associations between FFR periodicity and SIN perception might be mediated by right PAC [10, 11]. Our EEG findings suggest otherwise. Electrical FFRs failed to show a right hemisphere bias in cortical FFRs as suggested by MEG [10, 13, 25]. Rightward hemispheric bias in cortical responses to SIN have been widely reported in EEG studies [8, 28] and thus, were anticipated in the current study. Yet, we did not observe rightward lateralization of speech-FFRs, consistent with its bilateral symmetric scalp topography and deeper anatomical origin of midbrain responses [Fig 1b; 5, 6]. Instead, subcortical foci (AN, midbrain) were the dominant predictors of SIN perception as implied (but never confirmed) in previous FFR reports [e.g., 1, 7, 9, 22, 23, 27, 33]. In fact, despite being weakly observable, cortical FFRs here did not predict QuickSIN scores. Although we find moderate evidence favoring relations between *subcortical* FFR_EEG_ and SIN processing (Bayes factors = 3-4), we acknowledge limitations of our smaller sample size. Additional EEG studies on larger samples are needed to replicate and confirm present findings.

Discrepancies between neuroimaging studies likely reflect differences in the sensitivity of MEG vs. EEG for detecting deep vs. superficial neuronal currents. The bias of MEG to more superficial and tangential brain activity may account for the differential brain-behavior relations between FFRs and SIN perception observed in this (EEG) vs. previous (MEG) FFR studies.

Indeed, more recent MEG studies have begun to posit a brainstem locus when describing FFRs [17, 25, 37]. TMS-induced virtual lesions to auditory cortex also fail to eradicate FFRs [20], further suggesting a brainstem locus. Based on EEG and computational modeling, we have suggested the midbrain provides a 3× larger contribution to scalp FFRs (at F0) than cortex [6]. Similar conclusions were recently drawn by Ross, et al. [25], who further suggested cortical FFR sources are weaker when measured with EEG than with MEG. The disparity between MEG-and EEG-based FFRs is further bolstered by phase differences across the two modalities; for frequencies around 100 Hz, FFR_MEG_ group delays are later than those for FFR_EEG_, suggesting magnetic FFRs stem from later source(s) (i.e., cortex) than those measured via EEG [3]. We infer that the burden of SIN perception—with regards to the FFR_EEG_—is mainly on subcortical parts of the auditory neuroaxis and the degree to which neural periodicities capture and maintain the dynamic, time-frequency cues of speech.

## Acknowledgments

We thank Dr. Caitlin Price for comments on earlier versions of this manuscript. This work was supported by grants from the NIH/NIDCD (R01DC016267) awarded to G.M.B.

## Footnotes

1 Goodness of fit (GoF) = 1-RV, where RV is the residual variance computed as the sum of squares of the unexplained signal not accounted for by the source model.

2 One individual’s QuickSIN data was lost due to an error in data logging (EEG data were unaffected). This value was imputed using mean of the remaining participants’ QuickSIN scores.

3 Small peaks are visible near H2 for the PAC sources. These, however, were not significantly different from the EEG noise floor and thus, are likely some degree of cross-talk (leakage) of activity picked up from the midbrain BS source.

4 http://pcl.missouri.edu/bayesfactor

5 In light of potential problems with collinearity between bilateral sources we ran an additional GLME collapsing hemispheres for PAC and AN. BS (*p*=0.0015) and AN (*p*=0.0014) remained significant predictors of QuickSIN scores, whereas PAC was not (*p*=0.21).

## Supplementary Material

### SI Methods

#### Source latency analysis

We measured the latency of each FFR source to verify anatomical plausibility of our dipole model and estimate the propagation delay between successive auditory stages [3, 6]. Only clean responses were considered in this analysis given their more favorable SNR. The time lag between pairs of nuclei was estimated by cross-correlating their dipole source waveforms [4, 6, 9]. This provided a running correlation as a function of lag between each regional pair. The lag within a search window (1-11 ms) producing the maximum cross-correlation was taken as the latency shift between sources. This constraint was necessary given the periodic nature of the FFR and the largest pitch period of our stimulus (i.e., 88 Hz F0 ∼ 11 ms period) which yielded a periodic cross-correlation function. Values near the lower bound (1 ms) indicate poor separation of sources [4, 6]. The cumulative delay between consecutive FFR sources was used to confirm latencies fell within a plausible range of transmission time between auditory structures [4, 6].

Inter-regional source delays are shown in Figure S1. The lag between AN and IC was 5.1 ± 2.4 ms, consistent with the central transmission time from the cochlea to midbrain estimated via the click ABR I-V interval [7, 8, 10–12]. The cumulative lag between IC and PAC was 9.4 ± 4.1 ms, corresponding to an inter-region transmission time of ∼4.3 ms (=9.4 −5.3). Lastly, the cumulative lag between AN and PAC was 14.0 ± 3.5 ms, consistent with the earliest response onset observed in human depth electrode recordings from primary auditory cortex (∼13 ms) [17] and first spike-latencies in animal models [∼11-18 ms; 13, 26]. These delays are nearly identical to those observed in previous FFR_EEG_ [4] and FFR_MEG_ studies [6]. Few participants showed maximum cross-correlations at the lower bound of the search window (1 ms), indicating good spatiotemporal separation among the source signals of our model.

**Figure S1:**
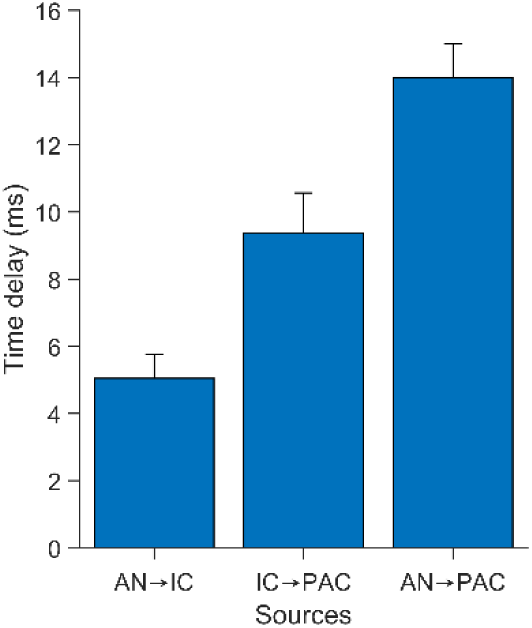
Cumulative lag between sources of the FFR. The propagation delay between successive sources was determined via cross-correlation of their dipole waveforms, constrained to latencies between one stimulus period (i.e., <11 ms).

#### Source sensitivity (spatial leakage)

Source models are prone to spatial leakage and cross-pickup of activity between distal sources. To assess the spatial specificity of our dipole model, we measured source sensitivity for each dipole location using BESA [for computational details, see 2]. Figure S2 shows the source sensitivity of each region in the 5-dipole model describing the FFR used in the main text. Source sensitivity reflects the spatial filter (sensitivity) of each source in the model to activity in surrounding brain regions. Sensitivity is defined as the fraction of power at a given source location mapped onto the selected point region. Sensitivity is independent of the recorded scalp signals and depends only on the source model leadfield (described by the sensor configuration, head model, and a regularization constant [1%]). While some mixing is unavoidable due to volume conduction, note the good spatial filtering of cortical (PAC) from subcortical (BS, AN) sources. In particular, the BS source does not “see” the PAC foci and vice versa, PAC sources are insensitive to midbrain BS activity. For a comparable analysis of spatial leakage for MEG FFRs, see [16]. This spatial isolation along with our response delay analysis (Fig. S1) indicate good spatiotemporal separation among the source signals of our model.

**Figure S2:**
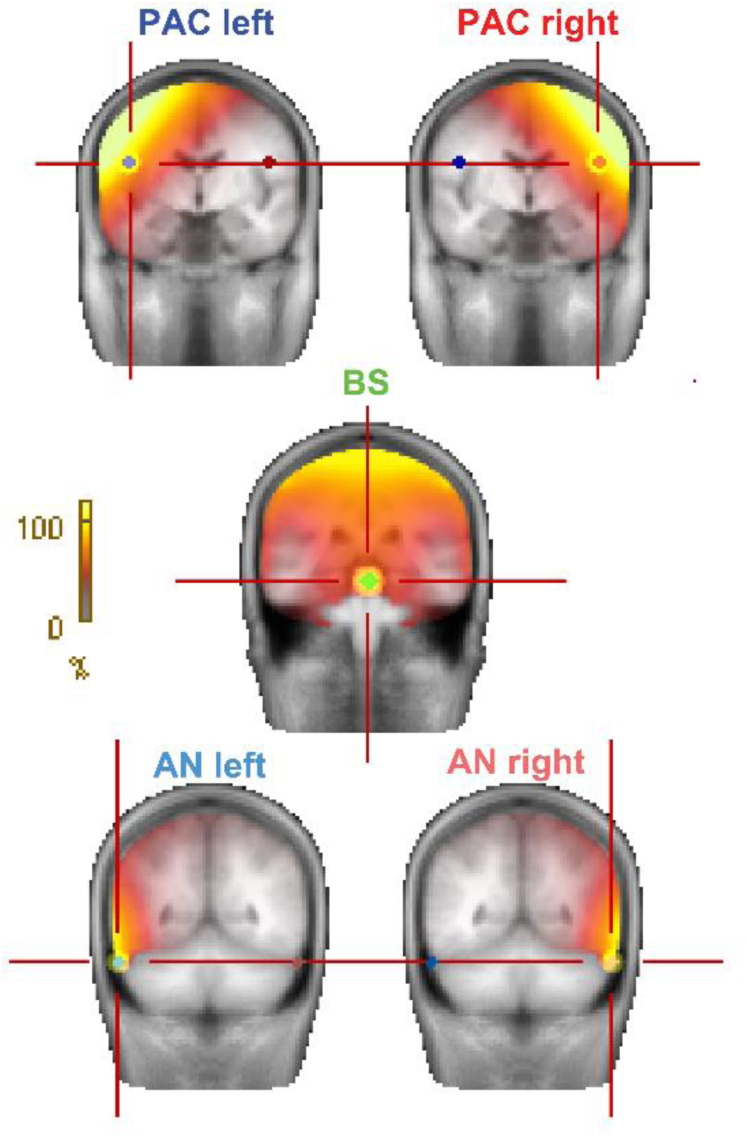
Source sensitivity (spatial filters) for the 5-dipole model describing the FFR (see Fig. 2, main text). Hotter colors denote a larger fraction of power at a given location that is mapped onto the selected source. Note the good spatial filtering and insensitivity of cortical to subcortical sources and vice versa.

### SI Discussion

Our neuroimaging data establish that the relative contribution of sources to the scalp FFR_EEG_ varies critically across levels of the auditory neuroaxis. In agreement with prior source imaging work [4], we found a gradient in contributions to the response, with the BS (midbrain) forming the largest component of the FFR_EEG_ (i.e., BS > AN > PAC). Our findings converge with human and animal studies suggesting subcortex (e.g., inferior colliculus and lower structures) as the primary source(s) of the FFR within the 20-200 Hz frequency bandwidth [4, 15, 20–22]. Our data lead us to reiterate the critical point that structures producing FFRs critically vary in both a level-(i.e., stage of brain) and stimulus-(i.e., frequency) dependent manner.

Diminishing FFR strength from brainstem to cortex is broadly consistent with the known phase-locking tolerance of nuclei along the ascending auditory pathway [14, 24–26]. Physiologically, reduced phase-locking could be due to increased synaptic jitter, organizational complexities of the pathway including (re)convergence, increasing time constants of neuronal membranes, and/or different ionic channels that cause a deterioration in temporal processing in more central neurons [23]. Collectively, these processes act as a low pass filter resulting in each anatomical division having a reduced upper limit of temporal precision. Indeed, phase-locking in auditory nerve persists up to >4-5 kHz [14], whereas the upper limit in midbrain neurons is ∼1100-1200 Hz [3, 18]. In stark contrast, locking in auditory cortical neurons among several species (monkeys, guinea pigs) is generally not observed above ∼100-120 Hz [1, 5, 14, 25]— even lower by some accounts [< 50 Hz; 19]. Our EEG data therefore agree with prior animal studies by showing limited phase-locked FFR activity in cortex up to a nominal frequency of 120 Hz. Nevertheless, given the reduced nature of these responses we are left to reaffirm that cortical contributions to the speech-FFR, when weakly present, are largely band-limited to the F0 component of low-pitch stimuli (e.g., male speech) [4].

